# A Novel Artificial Intelligence (AI) Method to Classify and Predict the Progression of Alzheimer’s Disease

**DOI:** 10.1101/2024.06.03.597177

**Authors:** Md Mehedi Hasan, Senjuti Rahman, Harshit Parmar, Suman K. Chowdhury

## Abstract

**Purpose:** The objective of this study was to develop a novel AI-ensembled network based on the most important features and affected brain regions to accurately classify and exhibit the pattern of progression of the stages of Cognitive Impairment (CI).

**Methods:** We proposed a novel ensembled architecture, 3D ResNet-18 - RF (Random Forest), and used this network to categorize the stages of Alzheimer’s disease (AD). The residual unit (blocks of ResNet) was introduced to the 3D Convolutional Neural network (CNN) to solve the degradation problem. It was considered an innovative strategy since the combination with fine-tuning resulted in higher accuracy. This network was trained on selected features and affected brain regions. The structured magnetic resonance images (MRI) were collected from the Alzheimer’s Disease Neuroimaging Initiative (ADNI) dataset, and the random forest was used for determining the importance of the features and affected regions from the parcellated 170 regions of interest (ROIs) using Atlas, automated anatomical labeling 3(AAL-3). This framework classified five categories of AD and detected the progression pattern.

**Results:** The proposed network showed promising results with a 66% F-1 score, 76% sensitivity, and 93.5% specificity, which outperformed the performance of conventional methods for categorizing five categories. Ventral Posterolateral and Pulvinar lateral regions were the regions most affected, indicating the progression from early MCI to AD. The five-fold validation accuracy for the developed model was 60.02%.

**Conclusion:** The results showed that the gray matter to white matter ratio was the most significant feature, which also accurately predicted the progression pattern. The performance metrics fluctuated with different hyperparameters, but they never exceeded 0.05% of the estimated results, indicating the validity and originality of the suggested methodology.

## Introduction

Alzheimer’s disease (AD) is a progressive neurodegenerative disease resulting in brain cell death and brain tissue shrinkage [1]. Adverse effects that people with AD progressively experience include breathing difficulties, heart failure, and, ultimately, death [2]. It is also the most prevalent form of dementia, which collectively indicates various forms of cognitive impairments (CI), such as memory loss, communication difficulty, and decline in thinking and reasoning skills [3]. According to the 2020 World Health Organization (WHO) report [6], approximately 55 million individuals have dementia and AD worldwide, and this number is predicted to almost double every 20 years, reaching 78 million in 2030 and 139 million by 2050 [6]. In the United States alone, about 5.8 million people aged 65 or older were living with AD in 2020 [7]. Approximately $321 billion was spent on the long-term medical care of Alzheimer’s patients in 2022, demonstrating the significant economic impact that the disease had on society [8].

The pathophysiological process of AD begins years before its clinical symptoms appear. The mild cognitive impairment (MCI) condition is considered a risk state that later progresses to AD [4]. In research and clinical settings, the progression of MCI is divided into two stages: early MCI (EMCI) and late MCI (LMCI). This classification distinguishes between individuals at the initial stage of clinical symptoms (EMCI) and those at a later stage (LMCI), progressing directly towards the development of AD [5]. As there is a lack of effective cure for AD, early and accurate detection is crucial [9]. Identifying the early stages of Alzheimer’s Disease (AD), such as EMCI and MCI through imaging modalities, would facilitate more effective treatment strategies for clinicians [10].

Previous studies have reported various MCI-associated brain degenerations, such as shrinking of the hippocampus, significant atrophy in the medial temporal lobe, and loss of gray matter volume in the frontal and parietal areas. Many neuroimaging studies conducted over the past few decades have attempted to advance the understanding of how to use imaging technologies like computed tomography (CT), PET (positron emission tomography), and MRI to diagnose human brain neurodegenerative disorders, including AD [11, 12]. MRI is the most commonly employed imaging technique for predicting stages of AD[13]. In order to reveal hidden information and eliminate noise inherent in complex MRI sequences, the extraction of important features becomes imperative[14]. Numerous studies have leveraged these critical input features extracted from MRI data to optimize performance [15–18]. The most commonly used MRI features include measuring brain volume, cortical thickness, brain surface area, texture, and white matter lesion (WML) volume, etc.[19, 20]. Deepti Putcha et al. [21] introduced a new approach to measuring structural brain changes by using the gray matter to white matter ratio, which emerged as a promising feature for detecting progression from earlier stages to the most prevalent stage (i.e., AD). Bruno Dubois et al. [22] discussed the development of vivo features that shifted the diagnosis of AD from the later dementia stages of the disease towards the earlier stages. They showed structural MRI reveals atrophy, or the loss of volume of gray matter, which denotes neurodegeneration in individuals with AD. Bijen Khagi et al. [23] revealed that a decrease in gray matter (GM) and brain volume, as well as a minor increase in cerebral-spinal fluid (CSF), can be an effective feature for the detection of AD. They used whole-brain MRI images to extract the abnormality of the GM and CSF regions to categorize impairments. The methods for covering the brain to extract features would be broadly divided into two categories: entire brain volume-based [5, 24] and region of interest (ROI))-based [24, 25]. ROI-based techniques frequently utilize 3D volume and shape properties of the specific brain regions. An atlas is a map that partitions the brain into anatomically distinct regions or ROI. In recent research papers, automated anatomical labeling 3 (AAL3) atlas was utilized to divide the brain regions for feature extraction after the pre-processing steps [26, 27]. Analyzing the volume, shape, and connectivity of specific brain regions has provided valuable insights into memory disorders, neurodegenerative diseases, and cognitive processes [28], thereby facilitating the classification of different categories of impairments.

Artificial intelligence, in particular machine learning and deep learning, is continuing to evolve and increasingly play a unique role in aiding MRI analysis in classifying the stages of AD [28–30]. Deep learning methods have been applied to predict AD using features extracted from structural MRI data [25, 30–32]. One of the most used deep learning networks is the Convolutional Neural network (CNN), which was used for the classification and prediction of AD [33–36]. Convolutional Neural Networks had advantages (increasing the depth and complexity of the network improved the classification performance) and disadvantages (gradient vanishing or explosion)[37]. Since MRI scans are three-dimensional, 3D CNN was able to interpret such images directly while maintaining the three-dimensional spatial context—a critical aspect of comprehending the internal structure of the brain [38]. In order to extract complicated features from 3D data and improve model comprehensibility, 3D CNNs were used in the prediction of Alzheimer’s disease [39]. In CNN, the feature maps from different layers did not interact much. The emergence of ResNet was a landmark in the development of deep learning methods, which brought in a residual unit that linked the current layer to the previous layer. The residual unit was introduced to solve the degradation problem [40]. The skip connection allowed ResNet to become deeper and show better performance than conventional CNN [41].

Zhuqing Long et al. [27] proposed a 3D densely connected convolutional neural network with a connection-wise attention mechanism to learn the multi-level features of brain MR images for AD classification. They used MRI of 968 subjects from the ADNI database to discriminate (1) AD versus healthy subjects, (2) MCI converters versus healthy subjects, and (3) MCI converters versus non-converters. Their proposed method achieved 97.35% accuracy for distinguishing AD patients from healthy control,87.82% for MCI converters against healthy control, and 78.79% for MCI converters against non-converters. In the training process, they adjusted the parameters of the deep CNN model, including the number of layers, the size, and the number of kernels in each layer; nevertheless, network convergence was still challenging. Furthermore, the study didn’t consider the multi-classification among the categories. Guilherme Folego et al. [42] developed an end-to-end deep 3D CNN (namely, LeNet-5, VGG, GoogLeNet, and ResNet 18) for the multiclass AD biomarker identification task, using the whole image volume as input. It was composed of three main steps: brain extraction and normalization, 3D CNN processing, and domain adaptation. It classified subjects into AD, MCI, or CN groups. They utilized the ADNI dataset and reached 52.3% accuracy in the testing set. They found the degradation problem, where traditional models similar to VGG stopped improving performance after a certain number of layers and even started getting worse afterward. To overcome this problem, they proposed the residual function, which was the basic building block of a Residual Network (ResNet). In their approach, the training was stopped after 50 epochs without further improvement in average TPF (true positive rate) over the validation set. This inspired us to use the skipnet feature from ResNet in our proposed network. Bijen Khagi et al. [43] used the SPM tool to get a 3D image of gray matter and fed it into 3D CNN. Once trained, an untested MRI can be passed through CNN to determine whether it is a healthy control (HC) or Mild Cognitive Impairment (MCI) due to AD (mAD) or AD dementia (ADD). They collected a dataset from the Gwangju Alzheimer’s Disease and Related Dementia (GARD) center. The dataset consisted of 42 Alzheimer’s disease dementia (ADD), 42 HC, and 39 MCI due to AD (mAD). They achieved an accuracy of around 40% for mutual information. The detailed feature extraction and analysis were still under study, and the overall classification result was not very high, which was due to the use of limited training materials. Yusi Chen et al.[44] introduced an ensemble deep learning model for AD classification, which incorporated Soft-NMS(Non-Maximum Suppression) into the Faster R–CNN architecture to classify the three categories, i.e., 115 AD patients, 106 subjects with MCI, and 185 CN subjects. Using a validation accuracy of about 50% and fine-tuning the ADNI dataset, they were able to reach an accuracy of 84.37% for the 3-way classification. They did not consider the five-way classification and recommended using a larger dataset and key features and regions as identification of early biomarkers for future work. We considered their recommendations for the proposed methodology by using important features and regions. Only a few studies [45, 46] have addressed the classification of the four stages of AD. However, to our knowledge, no study has explored the utilization of a five-way categorization scheme, coupled with the implementation of selected features using an ensemble network and a 3D deep learning network, for effectively categorizing the cognitive impairment condition associated with Alzheimer’s disease.

The objective of this study was to develop: i) a first-of-its-kind ensembled AI network to categorize the five stages of CI, (ii) the novel ensembled AI network comprising of an RF model to select the most important brain regions and features, and a novel fine-tuned 3D CNN (ResNet-18) architecture to categorize each disease condition accurately using structural MRI data retrieved from the ADNI database. The efficacy of our model was validated using a five-fold cross-validation approach, as well as existing literature data from similar works.

## Materials and Methods

The methodological approach of this work was divided into five major steps, comprising of 3D MRI data retrieved from the ADNI database (available at http://adni.loni.usc.edu), pre-processing, feature extraction, development of an ensembled AI network to predict CI condition using a minimal set of features. The entire process of the suggested methodology has been thoroughly introduced in this portion of the paper.

### Description of the ADNI Dataset

In this study, we used the ADNI database, which most of the researchers used to categorize the stages of Alzheimer’s disease [47–49]. In this study, we assessed T1-weighted MRI data from 1000 participants, including 200 AD patients, 200 LMCI subjects (developed to AD within 18 months of diagnosis), 200 MCI subjects (It did not develop to AD within 5 years of diagnosis), 200 EMCI subjects (AD was not diagnosed, early indication of AD), and 200 cognitively normal controls (CN) who had not changed over the course of 3 years [50]. Information about each group’s demographics and cognitive test results is shown in Table 1. It included categories, gender, no. of categories, average age, average CDR (Clinical Dementia Rate) score, and average MMSE score (Mini-Mental State Examination) for each category(reduces with the progression of Alzheimer’s disease[51]). The existence or absence of dementia and the degree of it were assessed using the CDR [52]. A global CDR of 0 denotes no dementia, a CDR of 0.5 denotes EMCI or very mild dementia, and a CDR of 1, 2, or 3 denotes mild, moderate, or severe dementia, where higher values correspond to more severe stages[53].

**Table 1.** Demographic and cognitive examination scores of the retrieved ADNI data.

| Category | #Subjects | Gender | | Age<br>average ± SD | MMSE<br>average ± SD | CDR<br>average ± SD |
| --- | --- | --- | --- | --- | --- | --- |
|  |  | F | M |  |  |  |
| **AD** | 200 | 100 | 100 | 74.40±5.78 | 20.16±6.60 | 2.98±0.60 |
| **LMCI** | 200 | 100 | 100 | 73.78±7.29 | 25.82±2.98 | 1.97±0.61 |
| **MCI** | 200 | 100 | 100 | 69.65±7.18 | 27.87±4.63 | 0.99±0.22 |
| **EMCI** | 200 | 100 | 100 | 72.078±8.05 | 28.86±1.99 | 0.435±0.20 |
| **CN** | 200 | 100 | 100 | 76.735±3.40 | 29.93±1.79 | 0.06±0.19 |

### Data Pre-Processing

Data preparation, pre-processing, segmentation, and post-processing were the four key phases of the proposed methodology. As part of the MRI preprocessing, we used an atlas to perform preprocessing procedures such as brain extraction, bias field correction, noise reduction, image registration, normalization, segmentation, and parcellation of 170 areas. In this study, statistical parametric mapping (SPM12) software was employed to complete the pre-processing steps, as most of the recent papers applied SPM12 for the preprocessing of the dataset [39, 40]. The feature extraction process was then carried out. Finally, we minimized the number of outliers after the dataset visualization because improper handling of them might lead to low accuracy.

An MRI of the brain reveals tissue as well as the spinal cord, skull, cranium, and eye [54]. Skull stripping should be used to separate the brain tissue from the non-brain tissue to distinguish the voxels as brain or non-brain. The brain stem, cerebral spinal fluid (CSF), Gray matter (GM), and White Matter (WM) of the cerebral cortex, as well as subcortical structures and the cerebellum, were examples of brain voxels, whereas non-brain voxels included the scalp, matter, eyes, bones, dura, skin, muscles, and fat [55]. In this study, the non-brain voxels were stripped before normalization and segmentation.

With the use of bias field correction, image contrast caused by magnetic field inhomogeneity [56] could be adjusted. When the MRI was conducted at 0.5 T, the bias field was essentially nonexistent and dependent on the strength of the magnetic field. However, it was regarded as powerful and could affect the MRI analysis when the magnetic field strength was 1.5 T, 3 T, or greater. We implemented the bias field correction step as it could greatly affect the MRI analysis. Noise reduction was the method of reducing the locally variable Rician noise seen in an MRI [56]. For categorization applications utilizing deep learning, this was thought to be less crucial [57]. In this research, noise reduction has been implemented to improve outcomes.

Image registration, which had two varieties, inter and intra-patient image registration — was primarily used to convert the alignment of the images from spatial to common anatomical spaces [58]. The dimensions of each MRI after registration were 79×95×79. Following that, each image was scaled uniformly through normalization and segmentation based on input modalities, which was carried out per the analysis’s goal. The segmented gray matter, white matter, and CSF are shown in Figure 1. A comprehensive parcellation of parts of the brain based on spatially normalized T1 volumes is offered by the Automated Anatomical Labeling (AAL) atlas. For parcellating brain regions, AAL3, a widely used atlas [44], was applied to the images to create 170 region masks for each subject set of data.

**Fig. 1.**
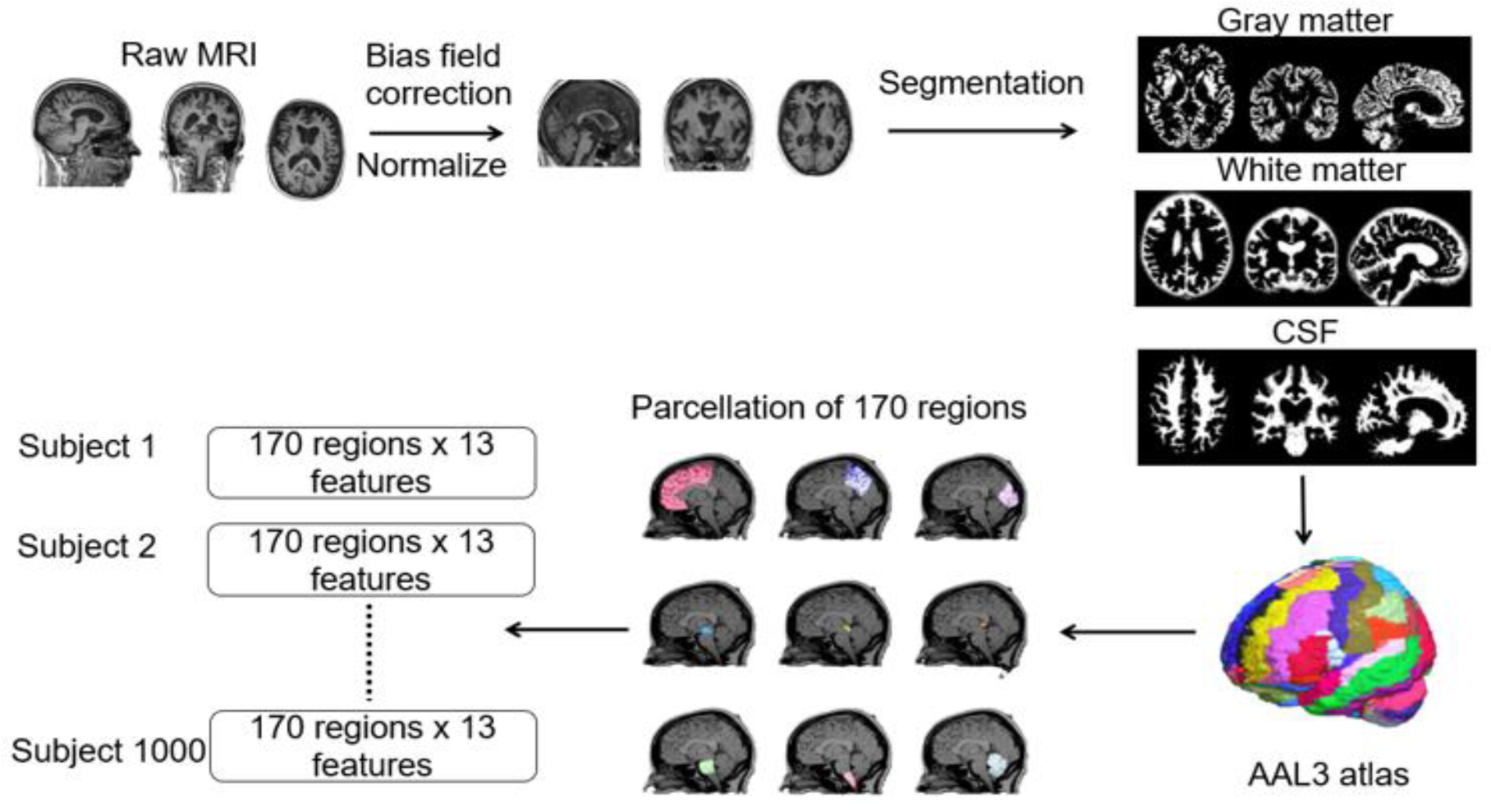
Pre-processing steps of MRI with respect to automated anatomical labelling atlas 3(AAL-3).

An outlier was an observation that was abnormally far from other values in a random sample taken from the population. Outliers can be found in a variety of ways. The interquartile range, or IQR, approach was one that was frequently applied [46]. We followed the IQR method and reduced the outliers as they could adversely affect the performance of the proposed method. We almost removed around 40 subjects from the whole dataset as they were identified as outliers.

#### Feature Extraction

After the preprocessing was accomplished, the preprocessed data were provided for the feature extraction. For the proposed methodology, feature extraction and ensembled model (RF-3D CNN) based classification were the two primary phases. Converting data into numerical values (feature extraction) was crucial for pattern recognition and visualization [10]. It was an important phase that helped the model identify more accurate results [58]. The feature extraction procedure extracted the features from images by selecting and combining variables into features to effectively reduce the amount of data. Following the creation of region masks using AAL 3 atlas, we counted the volumes of gray matter (GMV), white matter (WMV), and csf (CSFV) in each region and extracted their voxels. This was shown in equation (1).

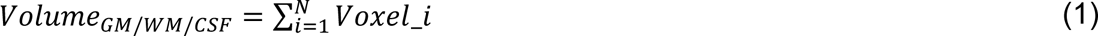

Where, voxel_i represented the volume of each voxel within the gray matter/white matter/cerebrospinal fluid region, and *N* was the total number of regions (ROIs). The same procedure was repeated for the five categories of AD. Then, for the first three features, the ratios of gray matter to csf (GCR), white matter to csf (WCR), and gray matter to white matter (GWR) were considered. GWR was considered as a novel feature by the previous studies where they considered a specific equation [21]. We used three equations to count the three ratios(shown in Table 2). Next, we divided the anatomical images into gray matter, white matter, and cerebrospinal fluid using the ground truth as the conventional Montreal Neurological Institute (MNI) template [59]. For this purpose, three ground truths—for gray matter, white matter, and CSF—had been used. As a result, three more features were obtained by counting the voxels, or volumes, from the ratios of white matter to white matter ground truth, CSF to CSF ground truth, and gray matter to gray matter ground truth. Several similarity metrics were applied in order to assess the overlap between the provided ground truth image and the anatomical brain MRI [60]. The Dice coefficient was the most well-known evaluation metric [61]. With this volume, the quality of two binary label masks could be compared. Let us define A as the mask produced by our proposed algorithm and B as the mask created by the standard MNI atlas. Equation (5) was then used to evaluate the dice overlap.

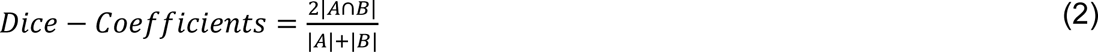

Where A = Gray matter/ White matter/ CSF volume, B = ground truth of gray matter/ White matter/ CSF volume

**Table 2.** ^1^Names and equations of the features.

| Feature Number | Name of the Features | Equations for the features |
| --- | --- | --- |
| Feature 1 | Gray matter to csf ratio (GCR) | $GCR = \frac{100 \times (GMV - CSFV)}{0.5(GMV + CSFV)}$ |
| Feature 2 | White matter to csf ratio (WCR) | $WCR = \frac{100 \times (WMV - CSFV)}{0.5(WMV + CSFV)}$ |
| Feature 3 | Gray matter to white matter ratio (GWR) | $GWR = \frac{100 \times (GMV - WMV)}{0.5(GMV + WMV)}$ |
| Feature 4 | Gray matter to gray matter ground truth ratio (GGR) | $GGR = \frac{GMV}{Ground\ truth\ of\ GMV(GGMV)}$ |

<sup>1</sup> GMV: volume of gray matter, WMV: volume of white matter, CSFV: volume of csf, N: total number of regions, Intensity<sub>i</sub> : the intensity value of the i-th region

|  |  |  |
| --- | --- | --- |
| Feature 5 | White matter to white matter ground truth ratio (WGR) | $WGR = \frac{WMV}{Ground\ truth\ of\ WMV(GWMV)}$ |
| Feature 6 | CSF to csf ground truth ratio (CGR) | $CGR = \frac{CSFV}{Ground\ truth\ of\ CSFV(GCSFV)}$ |
| Feature 7 | Dice coefficient of gray matter | $Dice - coefficient = \frac{2\vert GMV \cap GGMV\vert }{\vert GMV\vert + \vert GGMV\vert }$ |
| Feature 8 | Dice coefficient of white matter | $Dice - coefficient = \frac{2\vert WMV \cap GWMV\vert }{\vert WMV\vert + \vert GWMV\vert }$ |
| Feature 9 | Dice coefficient of csf | $Dice - coefficient = \frac{2\vert CSFV \cap GCSFV\vert }{\vert CSFV\vert + \vert GCSFV\vert }$ |
| Feature 10 | Average intensity | $Average\ intensity = \frac{1}{N} \sum_{i=1}^N Intensity\_i$ |
| Feature 11 | Surface area of gray matter | $Surface\ area = GMV - erosion\ of\ GMV$ |
| Feature 12 | Surface area of white matter | $Surface\ area = WMV - erosion\ of\ WMV$ |
| Feature 13 | Surface area of csf | $Surface\ area = CSFV - erosion\ of\ CSFV$ |

The overlap measure had values in the range of [0, 1], with 0 indicating no match between the two masks and 1 indicating that the two masks were identical. As a result, measurements of the three dice coefficients were made for each participant in each region. The anatomical image’s average intensity was then determined, making it the tenth feature. It was necessary to add the intensity of each mask for each region before dividing the result by the total number of voxels in order to obtain the average intensity of the anatomical image (the associated equation is shown in Table 2).

The erosion procedure had to be carried out to extract the final three features. The regions had to be scaled to do that, and the surface area was then calculated by subtracting from the main image using the AAL3 atlas. Next, the gray matter, white matter, and csf erosion ratios were calculated. The thirteen features extracted from the MRI were gray matter to CSF ratio, white matter to CSF ratio, gray matter to white matter ratio, gray matter to gray matter ground truth ratio, white matter to white matter ground truth ratio, CSF to CSF ground truth ratio, Dice coefficient between gray matter and gray matter ground truth, white matter and white matter ground truth, CSF and CSF ground truth, average intensity for gray matter, white matter and CSF and the surface area. The features and their associated equations are shown in Table 2. The visualization of extracting the 13 features from the brain atlas of 170 regions was shown in Figure 1.

### Development of Ensemble RF Network

The easiest way to describe Random Forest (RF), an ensemble machine learning technique, was as a mixture of tree predictors where each tree in the forest was dependent on the values of a random vector that was randomly sampled and had the same distribution for all the trees in the forest [62].

This algorithm yielded one of the best accuracies to date in many applications and offered significant advantages over previous methods in handling highly non-linear biological data, noise resilience, ease of tuning, and capacity for effective parallel processing [63–65]. Because of these advantages, RF was a great option for solving high-dimensional problems where a large number of features were frequently redundant.

An ensemble of decision trees was used to create a random forest, with each tree dividing the input space into a collection of rectangles in order to minimize the loss function [66]. Each variable was iterated over during the training phase of a single tree, and the variable and split point that yielded the best partition for the training data were chosen. The partition that resulted in the lowest total impurity between the two created subsets was the optimal one. Until a stopping requirement was satisfied—for example, a minimum number of points in the rectangle—the operation was repeated [67].

The significance of each individual input variable could be assessed using Random Forest classifiers [68]. We constructed a random forest model to assess the variable relevance as part of our work. This assessment first enabled us to figure out which factors were most important to include in the model during training. This variable’s relevance rating aids in the interpretation of each prediction result once the model has been trained.

In this study, random forests were used for features, and they mostly affected region selection. Three of the most important features were i) gray matter & white matter ratio(f3), ii) white matter to CSF ratio(f2), iii) gray matter to CSF ratio(f1) had been selected (with the threshold level = 1) from the thirteen features extracted above. And the most affected sixty regions had been selected to train the proposed model. The importance of features and most affected regions that we used in this paper were shown in the results section. The parameters of the random forest were selected, as shown in Table 3, for both the selection of important features and the most affected regions.

**Table 3.** Selected parameters for the random forest used in this study.

| Parameter Name | Description | Chosen Value |
| --- | --- | --- |
| n_estimators | The number of decision trees | 100 |
| max_features | The largest number of features to consider when branching | 5 |
| max_depth | The maximum depth of a single tree | 25 |
| min_samples_split | The minimum number of samples required to split an internal node | 6 |
| Min_samples_leaf | The minimum number of samples required to be at a leaf node. | 1 |

### Determination of Most Affected ROI Region

For determining the most affected regions from 170 regions, the normalized data for healthy control had been considered as the reference in this study. The process was divided into four parts. In the first step, we were required to determine the mean of the 170×13 (where 170 was the number of regions and 13 was the features) matrix for the 200 people belonging to the control category. In the second step, the standard deviation of the same matrix was calculated using the mean for 200 control patients. In the third step, the z-score was calculated, and the size of the matrix for the z-score was 170×13×200. The same procedure was followed to calculate the z-score of AD, LMCI, MCI, and EMCI patients while considering the mean and standard deviation of the control category. Then, Random Forest (described above) was used to determine the most affected regions.

### Formation of new 3D images with ROIs filled with selected features

After the features had been extracted for every subject (around 960 subjects due to removing the outliers), we took a whole brain atlas and filled each parcellated region with the corresponding feature value. We constructed three brain volumes using the three most significant features, which summed up to create one 4D matrix of size 79×9579×3. The dimensions of the 4D matrix were such that the 4^th^ dimension indicated that there would be three channels for three features. In a similar way, another 4D matrix (79×95×79×13) was made for 13 features. The 4D matrices would be the 3D convolutional neural network (CNN) input.

### 3D CNN to Categorize AD

#### Convolutional Neural Network

ResNet-18 is a variant of the ResNet architecture, a type of Convolutional Neural Network (CNN). The common approach for defining the CNN model was indicated by two main components: (i) a feature extraction part that consisted of a series of convolutional layers with max-pooling and an activation function, and (ii) a classifier part that typically consisted of fully connected layers [69]. An optimizer and a loss function were two crucial machine learning components that aid in enhancing a model’s performance. An optimizer could optimize the model’s parameters to minimize the loss function, which quantifies the discrepancy between a model’s expected and actual output. The key components of a CNN were the convolutional layer, RELU layer, pooling layer, fully connected layer, and SoftMax layer [70]. This convolutional layer’s primary function was to convolve the input image with a collection of learnable filters or kernels. The activation function was applied to the input element-by-element using the Rectified Linear Unit (ReLU) layer. In addition to assisting in reducing computational complexity, memory consumption, and parameter count, the pooling layer decreased the spatial size (width and height) of the input volume. The high-level filtered images from earlier layers were taken by a fully connected layer, which flattened and created a single vector that could be used as an input for a subsequent layer. The softmax layer was often used in the final layer of a neural network-based classifier. A vector of real numbers in the range (0, 1) that added up to one was created from an N-dimensional vector of real numbers using the softmax function.

#### Residual Blocks in ResNet-18 Architecture

CNNs had been modified to handle volumetric data, leading to the development of 3D CNNs (Tiwari et al., 2023). A more extensive examination was possible with 3D CNNs than with their 2D counterparts because they might collect both spatial and temporal or depth-related information in three-dimensional data (Ji et al., 2012). This had been particularly helpful in the field of medical imaging, where three-dimensional images were frequently employed in procedures like CT, MRI, and ultrasound examinations. They have been effective in accurately identifying and categorizing affected tissues faster than conventional methods (Hosny et al., 2018).

To address the vanishing gradient issue that raised during deep convolutional network training, the ResNet (residual network) was introduced in one study [71]. ResNet provided a skip connection that disregarded the non-linear transformations with an identity function and facilitated the training of networks that were significantly deeper than those previously employed. ResNet had the benefit of allowing gradients from earlier layers to flow straight through the identity function to later layers. Due to the advantages of using the skip connection of ResNet, we utilized this in the proposed methodology. As random forest was utilized for the feature and region selection, the proposed network combines ResNet-18 and random forest.

ResNet has several residual networks. The concept of residual blocks comes from solving the degradation problem. The accuracy becomes saturated as the network’s depth increases. Maybe all the complexities of our data have been sufficiently learned by the layers. Even if we try to increase the layers to get a better accuracy, then the degradation problem arises. A larger training error is caused by the additional layers. Degradation suggests that not all systems are similar or easily optimized. If a shallow network learns the features with 90% accuracy, the deep variation of the same model may achieve even lower accuracy. This was because optimization gets extremely difficult as network depth rises. To address the degradation issue, we have a setup known as skip-connections, which skips part of the levels in the architecture and feeds the previous layer’s output to the current location, helping in optimization[71].

The skip-connection skips two layers and outputs the input, ‘X’. This is referred to as a shortcut/skip connection because there are no extra parameters involved; we are simply delivering the previous information to the layer. F(X) is the mapping learned by the network, and F(X)+X is the new output/expected output(H(X)). And the residual was the difference between the expected output and the input. The normal skip connection was used when the dimension of X and the output of stacked layers were the same (residual block 1), whereas the linear projection on the skip connection is such that the dimension of X’ and the output of stacked layers were the same (residual block 2). Residual Block 1 contained a convolutional layer with a batch normalization layer followed by a RELU layer. Then, another convolutional layer, the batch normalization layer, was added sequentially. There were no additional layers in the skip connection, whereas, in residual block 2, one additional block (one convolutional layer + batch normalization layer) was added to the skip connection.

#### Detailed 3D CNN (ResNet-18) Network Architecture

The proposed ensembled methodology for this study is shown in Figure 2. The flow of feature selection and the input and output of the proposed methodologies was shown in Figure 2(a), where there were two ResNet-18 and the input is 4D matrix with ROIs filled with selected features (both thirteen and three most important features) were used to see the differences in results of performance parameters. The same random forest was also selected to determine the most affected regions (the most affected sixty regions) for the two Resnet-18 models. One methodology used the thirteen features and all the regions (parcellated 170 regions), whereas the second one utilized the random forest to select the important features and most affected regions. The proposed methodology for using three features is shown in Figure 2(b).

**Fig. 2.**
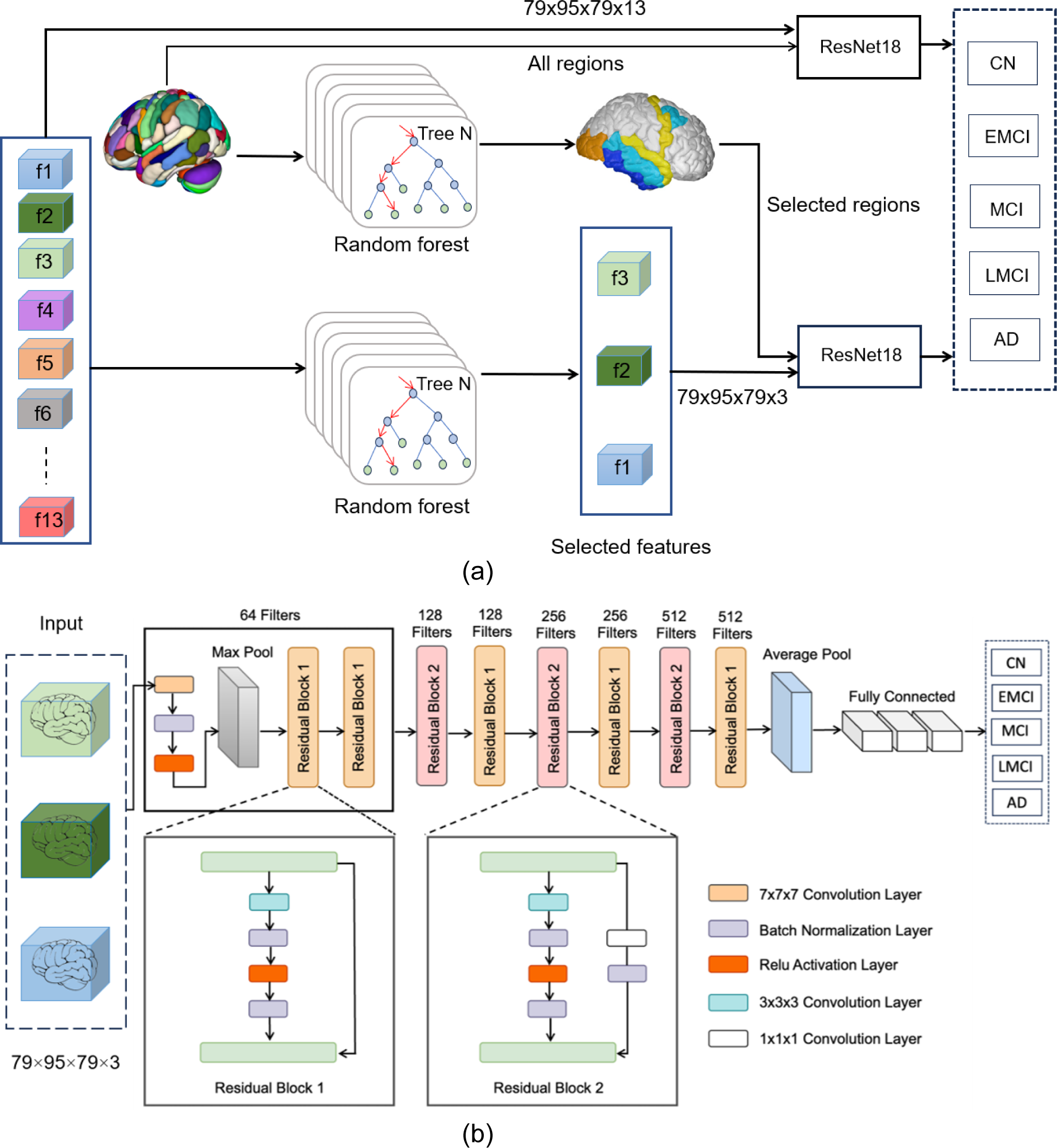
(a) The input and output of the proposed methodology, (b) The proposed ResNet-18 architecture.

The ResNet-18 architecture had a total of 18 learnable weight layers. The first layer was a 7×7×7 convolutional layer. Following the initial layer, 5 residual block 1 and 3 residual block 2. Each residual block 1 contains 2 weight layers (convolutional layers), and one skip connection (that connected the output of the second weight layer to the overall output using a RELU activation layer). Residual block 2 contained the same configuration as residual block 1 except the skip connection layer, which held one additional convolutional layer. The model architecture of the proposed method is shown in Table 4.

**Table 4.** ResNet-18 network architecture used in this study.

| Layer | Type | Size | Stride | Padding |
| --- | --- | --- | --- | --- |
| Input | 3D image input | 79×95×79×3 | -- | -- |
| Conv1 | 3D convolution | 40×48×40×64 | 2 2 2 | 3 3 3; 3 3 3 |
| Bn1 | Batch normalization | 40×48×40×64 | -- | -- |
| Relu1 | ReLU | 40×48×40×64 | -- | -- |
| Pool1 | 3D max pooling | 20×24×20×64 | 2 2 2 | 1 1 1; 1 1 1 |
| RS block 1 | Residual block 1 | 20×24×20×64 | 1 1 1 | 1 1 1; 1 1 1 |
| Relu2 | ReLU | 20×24×20×64 | -- | -- |
| RS block 1_2 | Residual block 1 | 20×24×20×64 | 1 1 1 | 1 1 1; 1 1 1 |
| RS block 2 | Residual block 2 | 10×12×10×128 | 2 2 2 | 1 1 1; 1 1 1 |
| RS block 1_3 | Residual block 1 | 10×12×10×128 | 1 1 1 | 1 1 1; 1 1 1 |
| RS block 2_1 | Residual block 2 | 5×6×5×256 | 2 2 2 | 1 1 1; 1 1 1 |
| RS block 1_4 | Residual block 1 | 5×6×5×256 | 1 1 1 | 1 1 1; 1 1 1 |
| RS block 2_3 | Residual block 2 | 3×3×3×512 | 2 2 2 | 1 1 1; 1 1 1 |
| RS block 1_5 | Residual block 1 | 3×3×3×512 | 1 1 1 | 1 1 1; 1 1 1 |
| Relu5 | ReLU | 3×3×3×512 | -- | -- |
| Pool5 | 3D average pooling | 1×1×1×512 | -- | -- |
| Fc1 | Fully Connected | 1×1×1×512 | -- | -- |
| Relu6 | ReLU | 1×1×1×512 | -- | -- |
| Dp1(70%) | Dropout | 1×1×1×512 |  |  |
| Fc2 | Fully Connected | 1×1×1×512 | -- | -- |
| Relu7 | ReLU | 1×1×1×512 | -- | -- |
| Dp2 (70%) | Dropout | 1×1×1×512 |  |  |
| Fc3 | Fully Connected | 1×1×1×5 | -- | -- |
| Prob | Softmax | 1×1×1×5 | -- | -- |
| Classification | Output (Cross Entropy) | 1×1×1×5 | -- | -- |

#### Training and Testing

We divided the whole dataset into training and testing datasets. Out of 960 subjects, the training, validation, and testing set had been portioned by 70:15:15 percent of the full dataset. We separated around 672 subjects’ data for training, and around 144 subjects were chosen randomly for validation purposes. The testing dataset comprised 144 subjects to test the proposed network. We also performed the five-fold cross-validation as it balances computational efficiency and model validation robustness[72]. There were a total of four training and testing stages for the cross-validation process, with all tasks of one participant being kept for the testing dataset in each case. The Adam optimizer and categorical cross-entropy loss function are commonly used in deep learning models, including those for Alzheimer’s disease research[73, 74]. We utilized the Adam optimizer and categorical cross-entropy loss function, which could contribute to these results by effectively training the model to classify patients based on their scans. We followed the flowchart shown in Figure 3 to train the proposed model. In the flowchart, we considered the threshold for validation of the five categories as equal to or greater than sixty percent, as most of the papers that considered four categories had validation accuracies around 60% [43, 75]. We trained the model until it had validation accuracy around the threshold value. We did not repeat the same procedure as in Figure 3 for two or three categories.

**Fig. 3.**
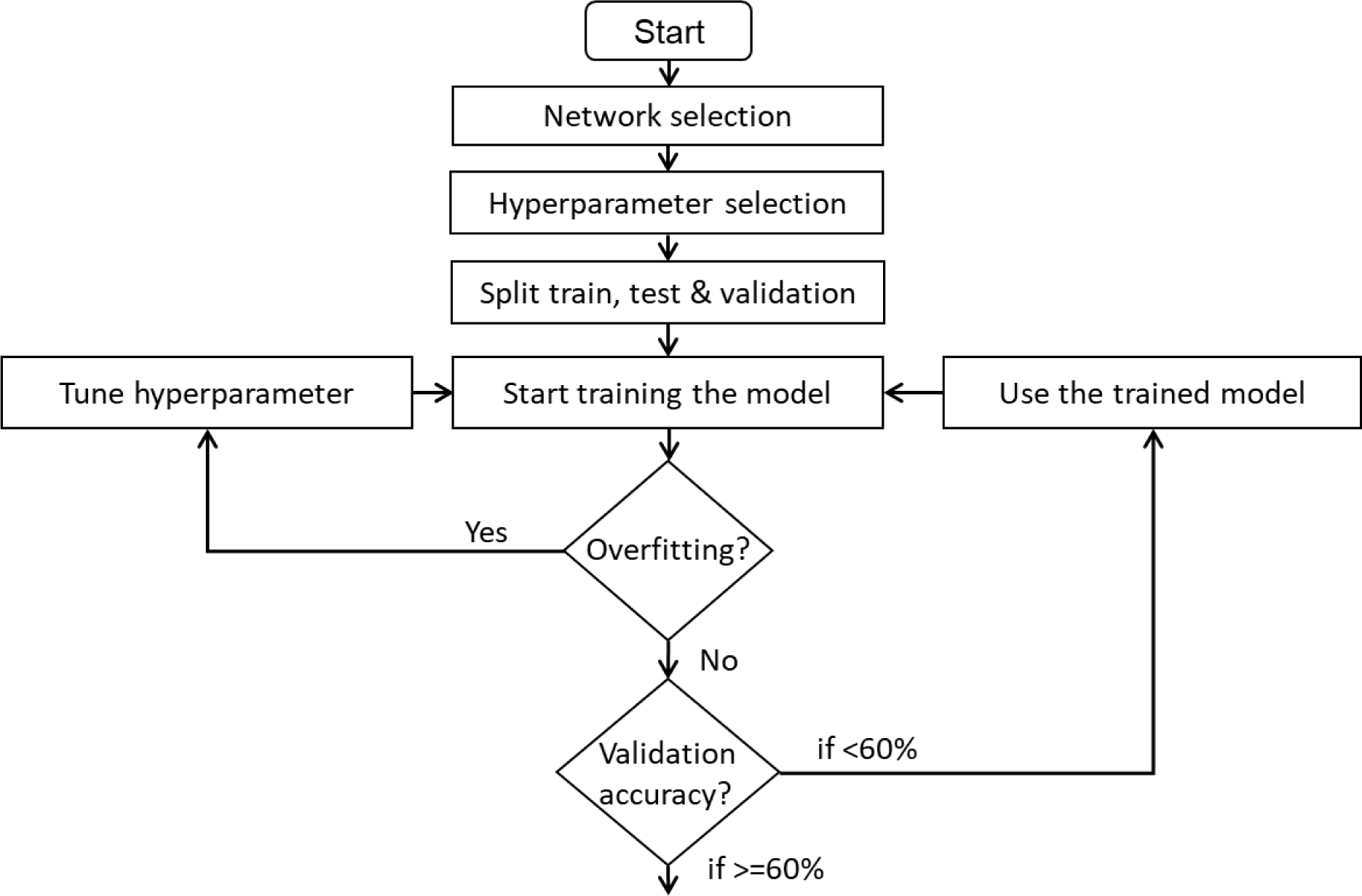
Method for training the proposed model.

#### Hyperparameters

We implemented the proposed 3D CNN (ResNet-18) on the thirteen features (all 170 regions) and, most importantly, three features most affected by sixty regions (depending on the importance categorization from the random forest). The highest accuracy was reported for the ResNet with three features sixty regions with 50 epochs. We trained our deep learning models on an Alienware Aurora R15(Windows 11 enterprise-64-bit) machine with 13th Gen Intel® Core i9 3.00 GHz CPUs (32 CPUs), 65536MB of memory, and a 64GB NVIDIA GeForce RTX 4090 GPU. The training converged in approximately 1.5(two categories) ±1.5 in total 3 hours for each model with a learning rate of 5e-05. The computation time depends on the features and regions. The combination of more features and regions led to more converging time, whereas the selected three features and sixty regions maintained the convergence within or less than 3 hours. The parameters for the highest accuracy are shown in Table 5.

**Table 5.**
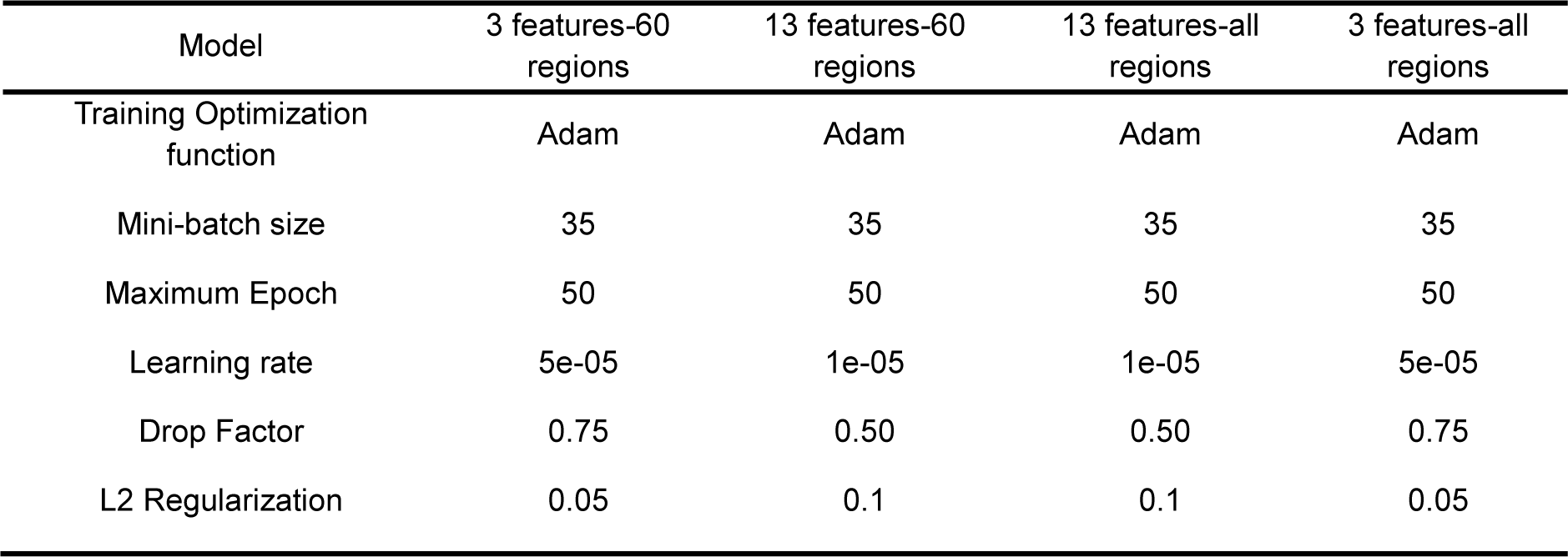
Description of hyperparameters and accuracy of the two approaches.

#### Validation

We used a five-fold cross-validation strategy to evaluate the classification performance. In all experiments, we split the data into five non-overlap folds, where one fold is used as the testing data, and the remaining four folds are used for training at each time. We repeat the whole process five times to avoid any possible bias caused by dataset partition. The final classification accuracy was reported by averaging the classification results from cross-validations.

## Results and Discussion

### Important Features and Most Affected Regions

Utilizing the ensembled network, ResNet-18, and random forest provided the performance parameters based on the significant attributes. Figure 4 displayed the feature importance graph with a threshold level of 1 arranged in descending order. From Figure 4, it was clear that three of the most important features were gray matter & white matter ratio(f3), white matter to CSF ratio (f2), and gray matter to CSF ratio(f1). To identify the most affected regions from 170 regions depending on the extracted features, colormap was used in the Matlab platform for the four categories: EMCI, MCI, LMCI, and AD. The most affected regions feature-wise, and common regions (basically the regions appearing in almost all the features) for each category were exhibited in Table 6. The anatomical description of the affected regions is also shown in Table 7. In Figure 5, the most affected sixty regions (in coronal, sagittal, axial, and render plane) were demonstrated for all the features of a patient of the MCI category, where blue indicates the least affected and red indicates highly affected regions.

**Fig. 4.**
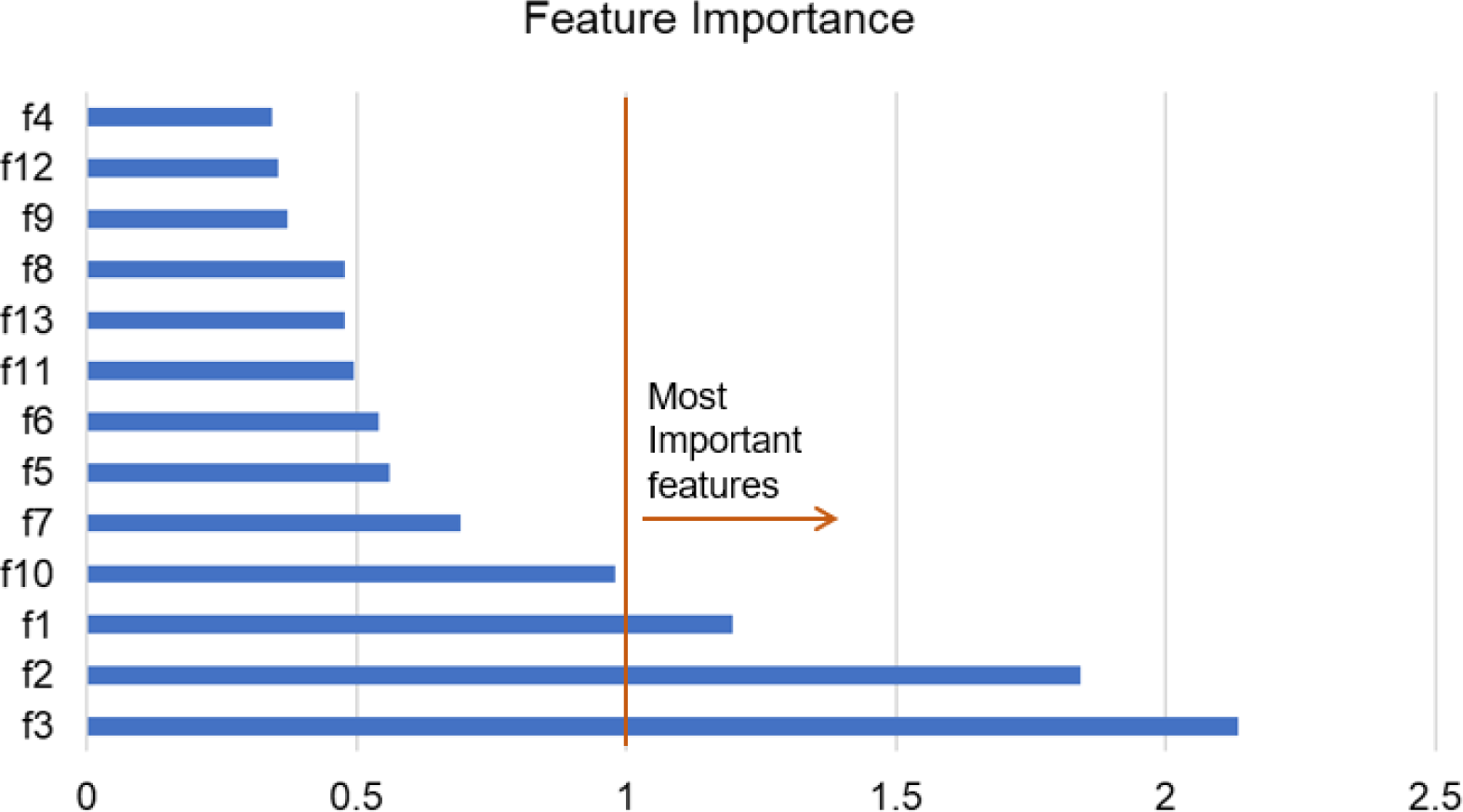
Feature importance in descending order.

**Fig. 5.**
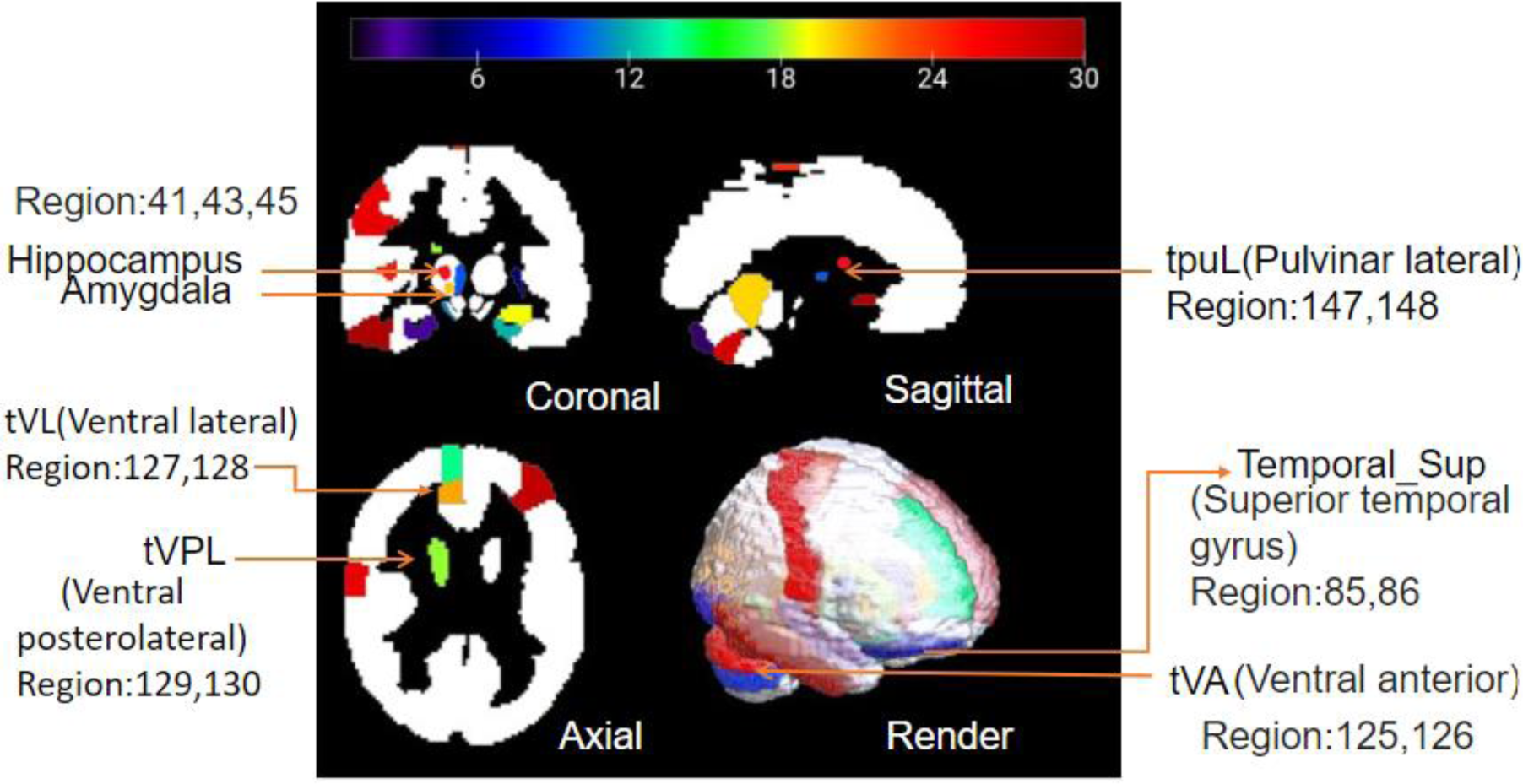
Mostly affected sixty regions for all categories depending on the thirteen features (for a subject of MCI Category).

**Table 6.**
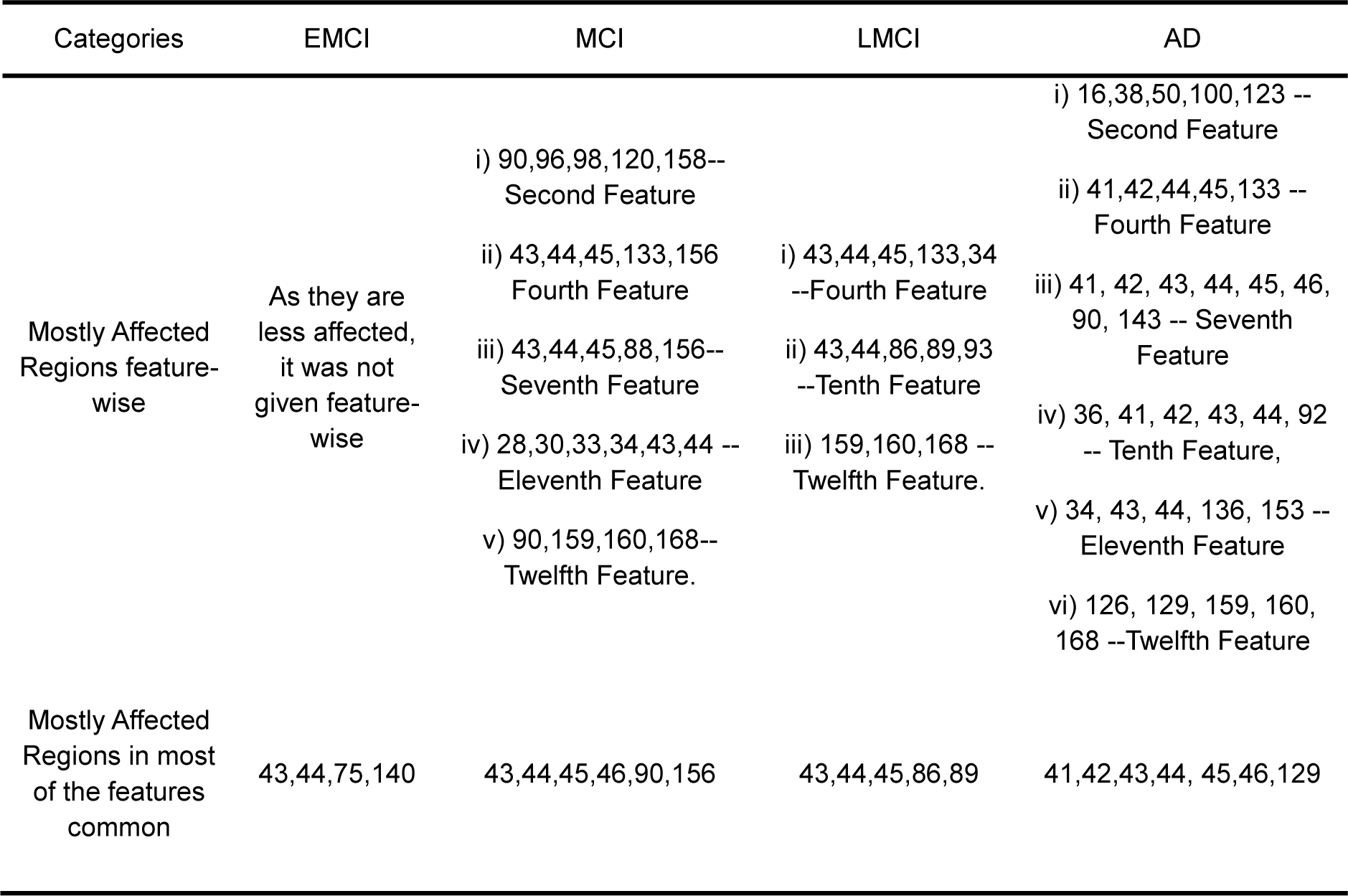
Most affected regions with respect to feature and commonly affected regions in all features.

**Table 7.** Anatomical description of the most affected regions for the categories[26].

| No. | Anatomical Description | Label |
| --- | --- | --- |
| 41,42 | Hippocampus left and right | Hippocampus |
| 43,44 | Left and right para-hippocampus | Para-Hippocampal |
| 45,46 | Left and right-amygdala | Amygdala |
| 75,76 | Caudate nucleus | Caudate |
| 79,80 | Lenticular nucleus, Pallidum | Pallidum |
| 85,86 | Superior temporal gyrus | Temporal_Sup |
| 87,88 | Temporal pole: superior temporal gyrus | Temporal_Pole_Sup |
| 89,90 | Middle temporal gyrus | Temporal_Mid |
| 121,122 | Thalamus, Anteroventral Nucleus | Thal_AV |
| 125,126 | Ventral anterior | Thal_VA |
| 127,128 | Ventral lateral | Thal_VL |
| 129,130 | Ventral posterolateral | Thal_VPL |
| 139,140 | Lateral geniculate | Thal_LGN |
| 147,148 | Pulvinar lateral | Thal_PuL |
| 155,156 | Anterior cingulate cortex, supracallosal | ACC_sup |

#### Ensembled Network Validation

The performance outcomes of the suggested models (including run 1, run 2, run 3, and run 4) and comparison between them are presented using sensitivity, specificity, and F1-score in Table 9. It was found that 3D CNN(ResNet-18) with three features and sixty regions model (run 1) had outperformed the same models for other combinations (run 2, run 3, and run 4). The training accuracies for the three features, sixty regions (run 1), and three features, all regions (run 4), were reported to be 99.99% and 99.96%, respectively. On the other hand, the training accuracies for run 2(thirteen features and sixty regions) and run 3(thirteen features and all regions) were 77.29% and 80.10%, respectively which was smaller than run 1 and run 4. The highest sensitivity, F-1 score, and specificity values were reported for all the runs included in Table 8.

**Table 8.** Performance parameters of the proposed approaches (for five categories)

| Run number | Regions and features | Sensitivity | F-1 score | Specificity |
| --- | --- | --- | --- | --- |
| Run 1 | 3 features – 60 regions | 0.760 | 0.660 | 0.935 |
| Run 2 | 13 features – 60 regions | 0.600 | 0.439 | 0.883 |
| Run 3 | 13 features – all regions | 0.630 | 0.470 | 0.840 |
| Run 4 | 3 features – all regions | 0.733 | 0.650 | 0.941 |

**Table 9.** Performance parameters of the proposed approaches (for two and three categories)

| Run number | Category | Sensitivity | F1 score | Specificity |
| --- | --- | --- | --- | --- |
| Run 1 | AD Vs. CN | 0.933 | 0.912 | 0.932 |
| Run 2 | EMCI vs. MCI | 0.866 | 0.787 | 0.860 |
| Run 3 | LMCI Vs. AD | 0.832 | 0.819 | 0.830 |
| Run 4 | MCI Vs. AD | 0.833 | 0.793 | 0.830 |
| Run 5 | MCI Vs. LMCI | 0.801 | 0.768 | 0.800 |
| Run 6 | MCI vs. CN | 0.702 | 0.700 | 0.702 |
| Run 7 | CN Vs. MCI Vs. AD | 0.800 | 0.727 | 0.900 |
| Run 8 | CN Vs. LMCI Vs. AD | 0.866 | 0.833 | 0.9667 |
| Run 9 | EMCI Vs. LMCI Vs. AD | 0.690 | 0.689 | 0.867 |
| Run 10 | CN Vs. MCI Vs. LMCI | 0.800 | 0.645 | 0.967 |

As seen from the above table, the combination of selected features (three features) and selected regions (sixty regions) gave the highest values for performance parameters. We also utilized the proposed model for two and three categories to validate the model based on the selected features and most affected regions. The performance parameters for the two categories are shown in Table 9. The highest training accuracy had been reported for AD Vs. CN (93.33%). The five-fold cross-validation gives a validation accuracy of about 60.02% for the proposed model for selected features and regions.

### Clinical Relevance

The most affected regions with respect to all thirteen features were shown in figure 6 as a histogram. The horizontal axis represents the 170 regions and vertical axis represents the mode of features (how frequent the features were responsible for the affected regions). The threshold was taken as 4 for the mode of features, which means if one specific region was not affected for at least four features, it was not shown in the figure. From the figure, it was clear that the most affected regions for all the categories were 43,79,85,87,122,125,126,127,128,129,147. The most affected sixty regions were considered for the classification of the five categories. The main purpose of this study was to show the progression of the categories. We used the mean z-score values for each category’s most affected sixty regions to show the progression of Alzheimer’s disease (shown in figure 7).

**Fig. 6.**
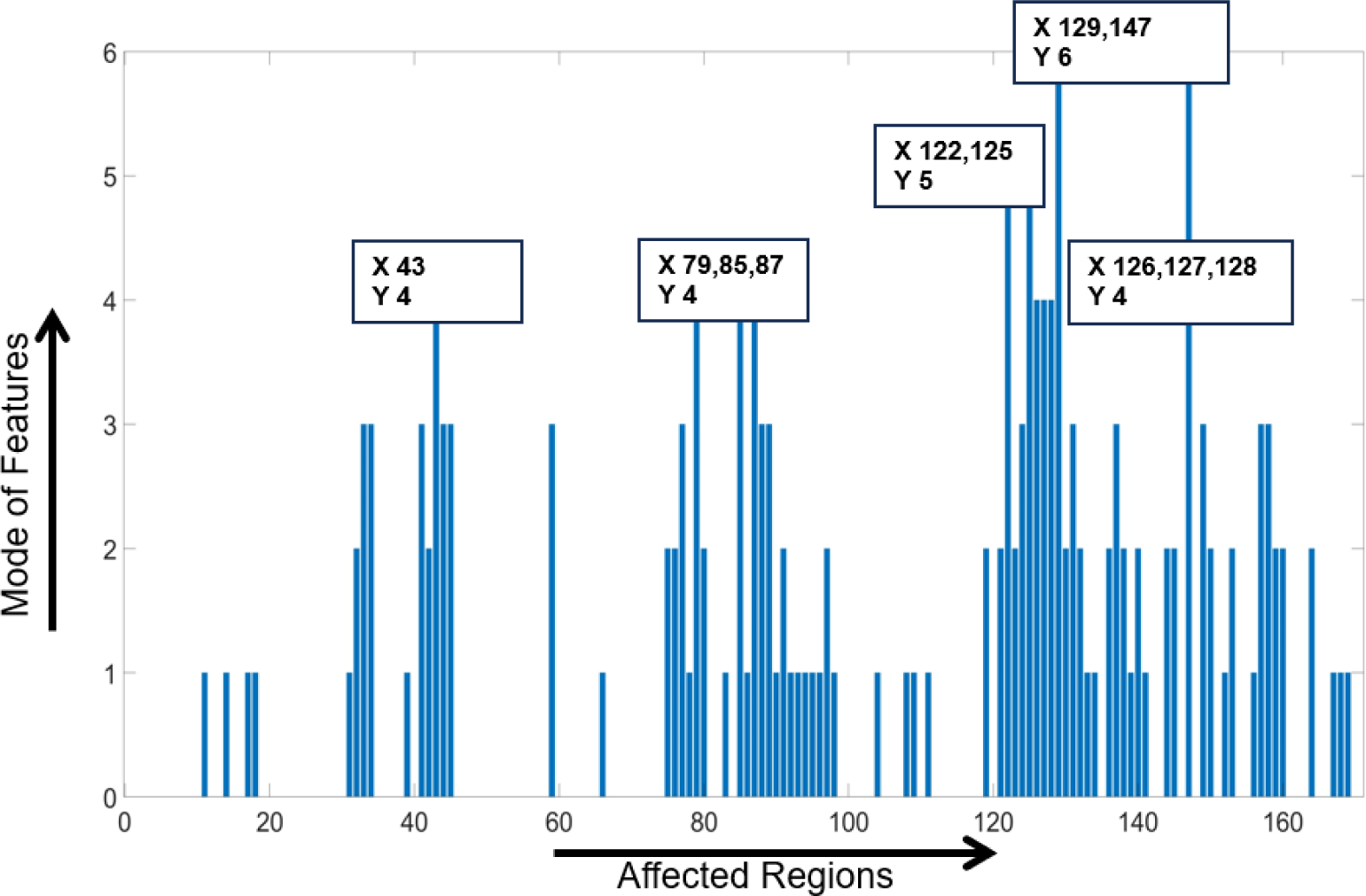
Most affected regions with respect to mode of thirteen features.

**Fig. 7.**
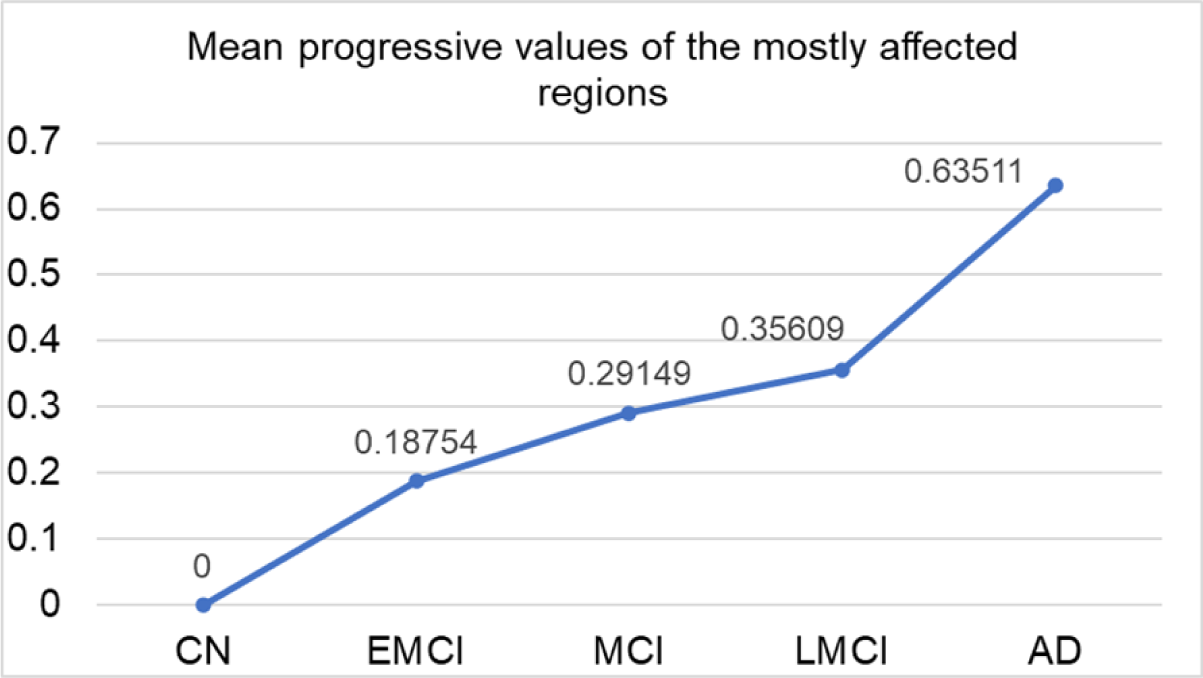
The Progression trend for the five categories based on mean Z-score value.

### Comparison with the Existing Work

Our research yielded better results than previous studies, as was evident when comparing it to a collection of existing literature. The comparison was made with similar methodology and ADNI dataset (most of them). The comparison is shown in table 10. From the table, it was evident that our ensembled network with selected features and regions provided best accuracy than the other network in the case of five categories and in some cases of two and three categories.

**Table 10.**
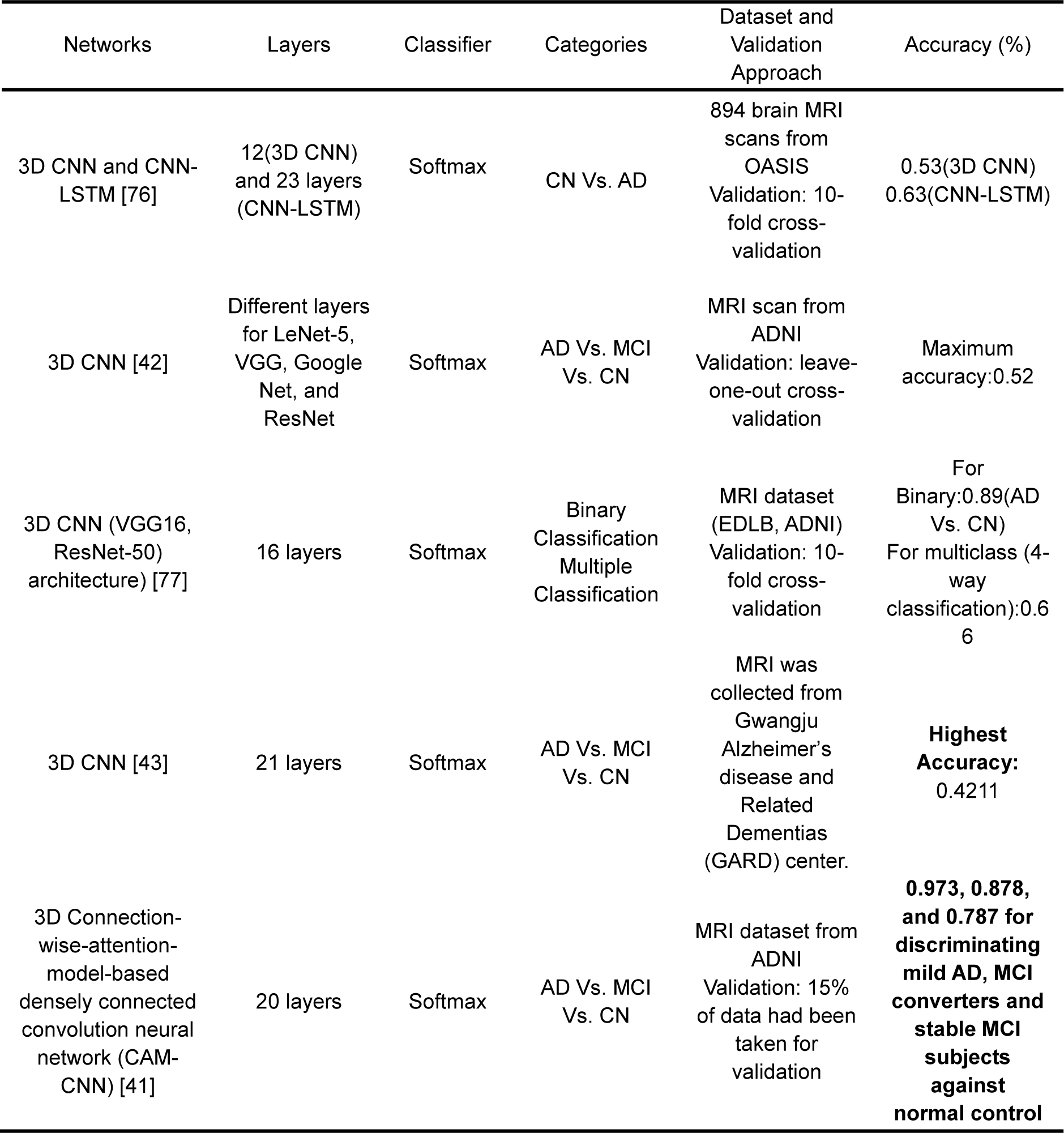
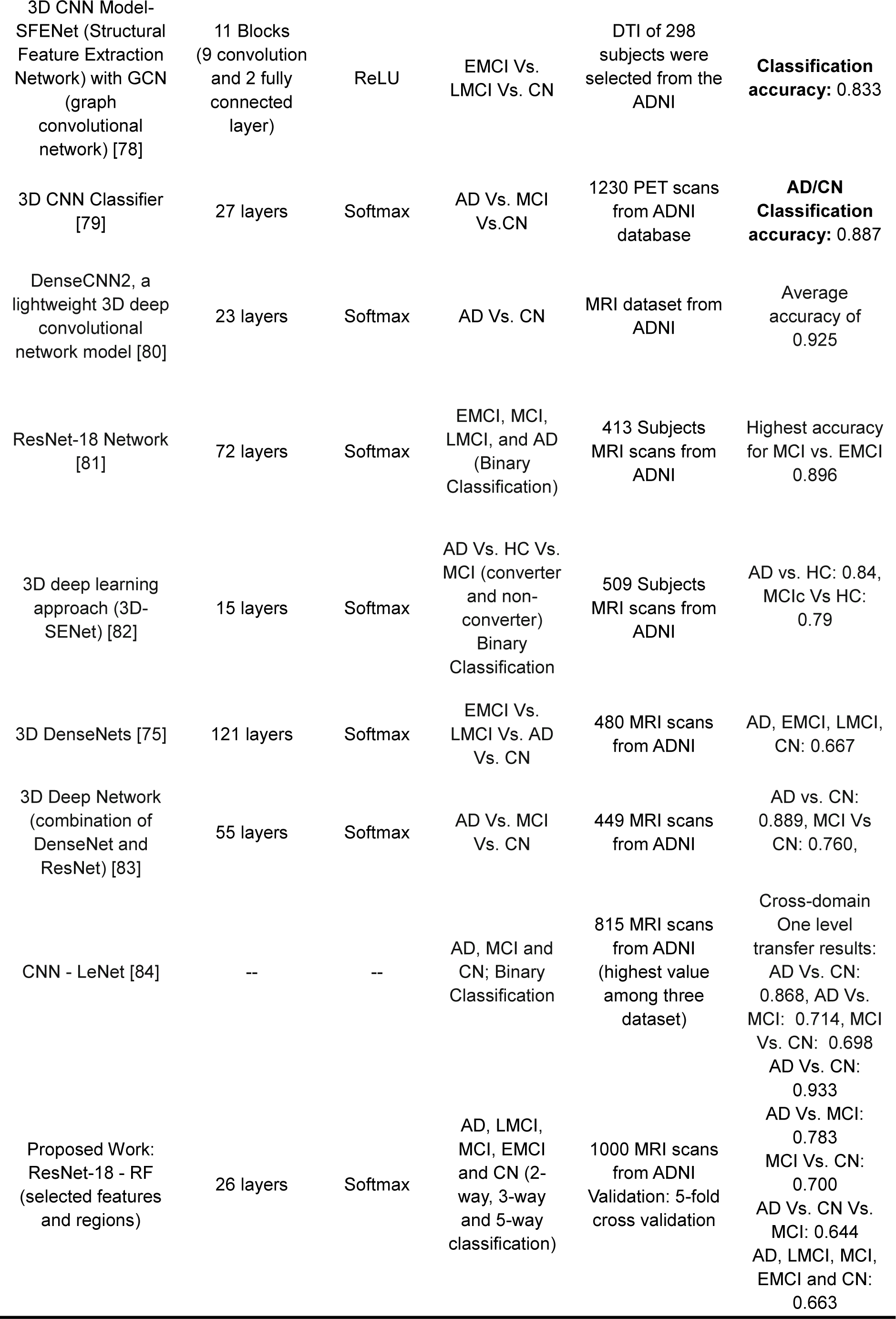
Comparison of the proposed model with the existing literature.

## Conclusion

In this study, the main focus was to classify the five categories of Alzheimer’s disease and predict the progression pattern, which would help clinicians detect the symptoms earlier so that they could increase the lifetime of the patients. From the findings, it was clear that the proposed ensembled network provides higher accuracies for the categorization of the stages of CI as it is considered the most important features and most affected regions. Furthermore, the extracted features were able to show the differences or progression patterns from EMCI to AD. It was also found that the hippocampus, amygdala, ventral posterolateral, pulvinar lateral regions, and GWR, WCR and GCR were the most affected regions, which gave significant progression patterns. From this study, we could build a relationship between the volumetric changes of the regions and progression according to the CI. As most of the studies considered two or three categories of Alzheimer’s disease, we also utilized the proposed ensembled network to validate our novel approach.

## Future Direction

Our proposed network gave a validation accuracy of nearly sixty percent, which could be improved by applying the transfer learning algorithm and by making the voxels as 2D slices. The total number of MRI samples was 1000, which could be increased to reduce the effects of overfitting. Different features (selection of different weights/coefficients and different combinations) and Atlases could be used to localize the brain regions and determine the progression rate for each combination of feature and Atlas. Furthermore, depending on the hippocampus and amygdala regions, the biomarker could be developed for different specific features.

## Acknowledgments

The work was not supported by any funding agencies.

## Footnotes

1 GMV: volume of gray matter, WMV: volume of white matter, CSFV: volume of csf, N: total number of regions, Intensity_i : the intensity value of the i-th region

## References

1. Muhammed Raees, P.C. and V. Thomas. Automated detection of Alzheimer’s Disease using Deep Learning in MRI. in Journal of Physics Conference Series. 2021.

2. Sharma, S., et al., A deep learning based convolutional neural network model with VGG16 feature extractor for the detection of Alzheimer Disease using MRI scans. Measurement: Sensors, 2022. 24: p. 100506.

3. Health, N.I.f. and C. Excellence, Dementia: assessment, management and support for people living with dementia and their carers. 2018: National Institute for Health and Clinical Excellence.

4. Mueller, S., et al., Functional and structural MR imaging in neuropsychiatric disorders, Part 1: imaging techniques and their application in mild cognitive impairment and Alzheimer disease. American journal of neuroradiology, 2012. 33(10): p. 1845–1850.

5. Klöppel, S., et al., Automatic classification of MR scans in Alzheimer’s disease. Brain, 2008. 131(3): p. 681–689.

6. World Health Organization. Dementia: a public health priority.

7. Alzheimer’s disease facts and figures. 2020.

8. 2022 Alzheimer’s disease facts and figures. Alzheimers Dement, 2022. 18(4): p. 700–789.

9. Kyaga, Z.N.V.G.S., The Early Diagnosis of Alzheimer’s Disease: A Patient-Centred Conversation with the Care Team. 2022.

10. Mosconi, L., et al., Early detection of Alzheimer’s disease using neuroimaging. Experimental gerontology, 2007. 42(1-2): p. 129–138.

11. Schwarz, C.G., Uses of human MR and PET imaging in research of neurodegenerative brain diseases. Neurotherapeutics, 2021. 18(2): p. 661–672.

12. Yen, C., C.-L. Lin, and M.-C. Chiang, Exploring the frontiers of neuroimaging: a review of recent advances in understanding brain functioning and disorders. Life, 2023. 13(7): p. 1472.

13. Kubi, P., et al. A Deep Learning-Based Technique to Determine Various Stages of Alzheimer’s Disease from 3D Brain MRI Images. in International Conference on Information Integration and Web Intelligence. 2023. Springer.

14. Chakraborty, D., et al., Deep learning-based feature extraction with MRI data in neuroimaging genetics for Alzheimer’s disease. Genes, 2023. 14(3): p. 626.

15. Diciotti, S., et al. Identification of mild Alzheimer’s disease through automated classification of structural MRI features. in 2012 Annual International Conference of the IEEE Engineering in Medicine and Biology Society. 2012. IEEE.

16. Zheng, Y., et al., Machine learning-based framework for differential diagnosis between vascular dementia and Alzheimer’s disease using structural MRI features. Frontiers in neurology, 2019. 10: p. 456891.

17. Sierra, H., et al., Confocal imaging-guided laser ablation of basal cell carcinomas: an ex vivo study. The Journal of investigative dermatology, 2015. 135(2): p. 612.

18. Lama, R.K., et al., Diagnosis of Alzheimer’s disease based on structural MRI images using a regularized extreme learning machine and PCA features. Journal of healthcare engineering, 2017. 2017.

19. Stephen, R., et al., Brain volumes and cortical thickness on MRI in the Finnish Geriatric Intervention Study to Prevent Cognitive Impairment and Disability (FINGER). Alzheimer’s research & therapy, 2019. 11: p. 1–10.

20. Seo, K., et al., Visualizing Alzheimer’s disease progression in low dimensional manifolds. Heliyon, 2019. 5(8).

21. Putcha, D., et al., Gray to white matter signal ratio as a novel biomarker of neurodegeneration in Alzheimer’s disease. NeuroImage: Clinical, 2023. 37: p. 103303.

22. Dubois, B., et al., Biomarkers in Alzheimer’s disease: role in early and differential diagnosis and recognition of atypical variants. Alzheimer’s Research & Therapy, 2023. 15(1): p. 175.

23. Khagi, B. and G.-R. Kwon, 3D CNN based Alzheimerâ€™ s diseases classification using segmented Grey matter extracted from whole-brain MRI. JOIV: International Journal on Informatics Visualization, 2021. 5(2): p. 200–205.

24. Long, X. and C. Wyatt. An automatic unsupervised classification of MR images in Alzheimer’s disease. in 2010 IEEE Computer Society Conference on Computer Vision and Pattern Recognition. 2010. IEEE.

25. Zhang, D., et al., Multimodal classification of Alzheimer’s disease and mild cognitive impairment. Neuroimage, 2011. 55(3): p. 856–867.

26. Rolls, E.T., et al., Automated anatomical labelling atlas 3. Neuroimage, 2020. 206: p. 116189.

27. Long, Z., et al., Identifying Alzheimer’s disease and mild cognitive impairment with atlas-based multi-modal metrics. Frontiers in Aging Neuroscience, 2023. 15.

28. Li, H., et al., A deep learning model for early prediction of Alzheimer’s disease dementia based on hippocampal magnetic resonance imaging data. Alzheimer’s & Dementia, 2019. 15(8): p. 1059–1070.

29. Islam, J. and Y. Zhang, Brain MRI analysis for Alzheimer’s disease diagnosis using an ensemble system of deep convolutional neural networks. Brain informatics, 2018. 5: p. 1–14.

30. Pan, D., et al. Early diagnosis of Alzheimer’s disease based on deep learning and GWAS. in Human Brain and Artificial Intelligence: First International Workshop, HBAI 2019, Held in Conjunction with IJCAI 2019, Macao, China, August 12, 2019, Revised Selected Papers 1. 2019. Springer.

31. Ebrahimighahnavieh, M.A., S. Luo, and R. Chiong, Deep learning to detect Alzheimer’s disease from neuroimaging: A systematic literature review. Computer methods and programs in biomedicine, 2020. 187: p. 105242.

32. Liu, S., et al., Multimodal neuroimaging feature learning for multiclass diagnosis of Alzheimer’s disease. IEEE transactions on biomedical engineering, 2014. 62(4): p. 1132–1140.

33. Alzubaidi, L., et al., Review of deep learning: Concepts, CNN architectures, challenges, applications, future directions. Journal of big Data, 2021. 8: p. 1–74.

34. Jain, R., et al., Convolutional neural network based Alzheimer’s disease classification from magnetic resonance brain images. Cognitive Systems Research, 2019. 57: p. 147–159.

35. Lin, W., et al., Convolutional neural networks-based MRI image analysis for the Alzheimer’s disease prediction from mild cognitive impairment. Frontiers in neuroscience, 2018. 12: p. 777.

36. Huang, Y., et al., Diagnosis of Alzheimer’s disease via multi-modality 3D convolutional neural network. Frontiers in neuroscience, 2019. 13: p. 509.

37. Cong, S. and Y. Zhou, A review of convolutional neural network architectures and their optimizations. Artificial Intelligence Review, 2023. 56(3): p. 1905–1969.

38. Dyrba, M., et al., Improving 3D convolutional neural network comprehensibility via interactive visualization of relevance maps: evaluation in Alzheimer’s disease. Alzheimer’s research & therapy, 2021. 13: p. 1–18.

39. Xu, X., et al., A review of the application of three-dimensional convolutional neural networks for the diagnosis of Alzheimer’s disease using neuroimaging. Reviews in the Neurosciences, 2023(0).

40. Hu, Y., H. Tang, and G. Pan, Spiking deep residual networks. IEEE Transactions on Neural Networks and Learning Systems, 2021.

41. Zhang, J., et al., A 3D densely connected convolution neural network with connection-wise attention mechanism for Alzheimer’s disease classification. Magnetic Resonance Imaging, 2021. 78: p. 119–126.

42. Folego, G., et al., Alzheimer’s disease detection through whole-brain 3D-CNN MRI. Frontiers in bioengineering and biotechnology, 2020. 8: p. 534592.

43. Khagi, B. and G.-R. Kwon, 3D CNN based Alzheimer’s diseases classification using segmented Grey matter extracted from whole-brain MRI. JOIV: International Journal on Informatics Visualization, 2021. 5(2): p. 200–205.

44. Chen, Y., et al., Automated Alzheimer’s disease classification using deep learning models with Soft-NMS and improved ResNet50 integration. Journal of Radiation Research and Applied Sciences, 2024. 17(1): p. 100782.

45. Pasnoori, N., T. Flores-Garcia, and B.D. Barkana, Histogram-based features track Alzheimer’s progression in brain MRI. Scientific Reports, 2024. 14(1): p. 257.

46. Raza, N., et al., Alzheimer disease classification through transfer learning approach. Diagnostics, 2023. 13(4): p. 801.

47. Diogo, V.S., et al., Early diagnosis of Alzheimer’s disease using machine learning: a multi-diagnostic, generalizable approach. Alzheimer’s Research & Therapy, 2022. 14(1): p. 107.

48. Rajesh Khanna, M., Multi-level classification of Alzheimer disease using DCNN and ensemble deep learning techniques. Signal, Image and Video Processing, 2023: p. 1–9.

49. Toga, A.W. and K.L. Crawford, The Alzheimer’s Disease Neuroimaging Initiative informatics core: a decade in review. Alzheimer’s & Dementia, 2015. 11(7): p. 832–839.

50. Wyman, B.T., et al., Standardization of analysis sets for reporting results from ADNI MRI data. Alzheimer’s & Dementia, 2013. 9(3): p. 332–337.

51. Henneges, C., et al., Describing the sequence of cognitive decline in Alzheimer’s disease patients: results from an observational study. Journal of Alzheimer’s Disease, 2016. 52(3): p. 1065–1080.

52. Morris, J.C., The Clinical Dementia Rating (CDR) current version and scoring rules. Neurology, 1993. 43(11): p. 2412–2412-a.

53. Tariot, P.N., et al., Relationships of change in Clinical Dementia Rating (CDR) on patient outcomes and probability of progression: observational analysis. Alzheimer’s Research & Therapy, 2024. 16(1): p. 36.

54. Duta, N. and M. Sonka, Segmentation and interpretation of MR brain images. An improved active shape model. IEEE transactions on medical imaging, 1998. 17(6): p. 1049–1062.

55. Rogowska, J., Overview and fundamentals of medical image segmentation. Handbook of medical imaging, processing and analysis, 2000: p. 69–85.

56. Vovk, U., F. Pernus, and B. Likar, A review of methods for correction of intensity inhomogeneity in MRI. IEEE transactions on medical imaging, 2007. 26(3): p. 405–421.

57. Coupé, P., et al., An optimized blockwise nonlocal means denoising filter for 3-D magnetic resonance images. IEEE transactions on medical imaging, 2008. 27(4): p. 425–441.

58. Klein, A., et al., Evaluation of 14 non-linear deformation algorithms applied to human brain MRI registration. Neuroimage, 2009. 46(3): p. 786–802.

59. Wu, Z., et al., Gray matter deterioration pattern during Alzheimer’s disease progression: a regions-of-interest based surface morphometry study. Frontiers in Aging Neuroscience, 2021. 13: p. 593898.

60. Carass, A., et al., Longitudinal multiple sclerosis lesion segmentation: resource and challenge. NeuroImage, 2017. 148: p. 77–102.

61. Dice, L.R., Measures of the amount of ecologic association between species. Ecology, 1945. 26(3): p. 297–302.

62. Breiman, L., Random forests. Machine learning, 2001. 45: p. 5–32.

63. De Bruyn, T., et al., Structure-based identification of OATP1B1/3 inhibitors. Molecular pharmacology, 2013. 83(6): p. 1257–1267.

64. Caruana, R. and A. Niculescu-Mizil. An empirical comparison of supervised learning algorithms. in Proceedings of the 23rd international conference on Machine learning. 2006.

65. Menze, B.H., et al., A comparison of random forest and its Gini importance with standard chemometric methods for the feature selection and classification of spectral data. BMC bioinformatics, 2009. 10: p. 1–16.

66. Moore, P., et al., Random forest prediction of Alzheimer’s disease using pairwise selection from time series data. PloS one, 2019. 14(2): p. e0211558.

67. Hastie, T., et al., The elements of statistical learning: data mining, inference, and prediction. Vol. 2. 2009: Springer.

68. Velazquez, M., Y. Lee, and A.s.D.N. Initiative, Random forest model for feature-based Alzheimer’s disease conversion prediction from early mild cognitive impairment subjects. Plos one, 2021. 16(4): p. e0244773.

69. Khoshdeli, M., R. Cong, and B. Parvin. Detection of nuclei in H&E stained sections using convolutional neural networks. in 2017 IEEE EMBS International Conference on Biomedical & Health Informatics (BHI). 2017. IEEE.

70. Phung, V.H. and E.J. Rhee, A high-accuracy model average ensemble of convolutional neural networks for classification of cloud image patches on small datasets. Applied Sciences, 2019. 9(21): p. 4500.

71. He, K., et al. Deep residual learning for image recognition. in Proceedings of the IEEE conference on computer vision and pattern recognition. 2016.

72. Zhang, F., et al., Accelerating Hyperparameter Tuning in Machine Learning for Alzheimer’s Disease With High Performance Computing. Frontiers in Artificial Intelligence, 2021. 4: p. 798962.

73. El-Latif, A.A.A., et al., Accurate Detection of Alzheimer’s Disease Using Lightweight Deep Learning Model on MRI Data. Diagnostics, 2023. 13(7): p. 1216.

74. AbdulAzeem, Y., W.M. Bahgat, and M. Badawy, A CNN based framework for classification of Alzheimer’s disease. Neural Computing and Applications, 2021. 33: p. 10415–10428.

75. Ruiz, J., et al. 3D DenseNet ensemble in 4-way classification of Alzheimer’s disease. in Brain Informatics: 13th International Conference, BI 2020, Padua, Italy, September 19, 2020, Proceedings 13. 2020. Springer.

76. Liu, W., J. Zhang, and Y. Zhao. A Comparison of Deep Learning and Traditional Machine Learning Approaches in Detecting Cognitive Impairment Using MRI Scans. in 2022 IEEE 46th Annual Computers, Software, and Applications Conference (COMPSAC). 2022. IEEE.

77. Soliman, A., et al., Adopting transfer learning for neuroimaging: a comparative analysis with a custom 3D convolution neural network model. BMC medical informatics and decision making, 2022. 22(Suppl 6): p. 318.

78. Kong, H., et al. Adversarial Learning Based Structural Brain-Network Generative Model for Analyzing Mild Cognitive Impairment. in Chinese Conference on Pattern Recognition and Computer Vision (PRCV). 2022. Springer.

79. Islam, J. and Y. Zhang, Understanding 3D CNN behavior for Alzheimer’s disease diagnosis from brain PET scan. arXiv preprint arXiv:1912.04563, 2019.

80. Katabathula, S., Q. Wang, and R. Xu, Predict Alzheimer’s disease using hippocampus MRI data: a lightweight 3D deep convolutional network model with visual and global shape representations. Alzheimer’s Research & Therapy, 2021. 13(1): p. 1–9.

81. Nicholas, P.J., et al., Using a ResNet-18 Network to Detect Features of Alzheimer’s Disease on Functional Magnetic Resonance Imaging: A Failed Replication. Comment on Odusami et al. Analysis of Features of Alzheimer’s Disease: Detection of Early Stage from Functional Brain Changes in Magnetic Resonance Images Using a Fine-tuned ResNet18 Network. Diagnostics 2021, 11, 1071. Diagnostics, 2022. 12(5): p. 1094.

82. Pan, D., et al., Early detection of Alzheimer’s disease using magnetic resonance imaging: a novel approach combining convolutional neural networks and ensemble learning. Frontiers in neuroscience, 2020. 14: p. 259.

83. Liu, M., et al., A multi-model deep convolutional neural network for automatic hippocampus segmentation and classification in Alzheimer’s disease. Neuroimage, 2020. 208: p. 116459.

84. Aderghal, K., et al., Improving Alzheimer’s stage categorization with Convolutional Neural Network using transfer learning and different magnetic resonance imaging modalities. Heliyon, 2020. 6(12).

